# Scaling ecological niches from individuals to populations and beyond

**DOI:** 10.1101/2024.12.06.627157

**Authors:** Muyang Lu, Scott Yanco, Ben Carlson, Kevin Winner, Jeremy Cohen, Diego Ellis-Soto, Shubhi Sharma, Will Rogers, Walter Jetz

## Abstract

The niche is a key concept that unifies ecology and evolutionary biology. However, empirical and theoretical treatments of the niche are mostly performed at the species level, neglecting individuals as important units of ecological and evolutionary processes. So far, a formal mathematical link between individual-level niches and higher organismal-level niches has been lacking, hampering the unification of ecological theories and more accurate forecasts of biodiversity change. To fill in this gap, we propose a bottom-up approach to derive population and higher organismal-level niches from individual niches. We demonstrate the power of our framework by showing that 1) the statistical properties of higher organismal-level niches (e.g. niche breadth, skewness etc.) can be partitioned into individual contributions; 2) the species-level niche shifts can be estimated by tracing the responses of individuals. Our method paves the way for a unifying niche theory and enables mechanistic assessments of organism-environment relationships across organismal scales.

## Introduction

The niche is one of the few unifying concepts in ecology and evolutionary biology across spatial, temporal, and organismal scales (1–3). The idea that every individual, population, and species should have a favorable set of environments (abiotic or biotic) allowing its persistence has been deeply ingrained in ecologists’ minds since Grinnell and Hutchinson (4–6). The niche concept underlies almost every explanation of and forecast for biodiversity change: from the accumulation of species richness over geologic time scales (7–9), the maintenance of diversity through coexistence mechanisms and ecological drift (10–13), to the predicted sudden collapse of ecosystems under global change (14, 15).

However, despite the nominal conceptual unification of the niche (16, 17), its empirical estimation with environmental data is almost exclusively focused on the species level (18–20). Such a species-centric view omits a simple but fundamental fact: the species niche, as well as the niches of higher-taxonomic levels (e.g. genus and families), is an emergent property of the constituent individuals (21–24). The conventional species-centric view overlooks the role of individuals as the fundamental units of ecological and evolutionary processes (25). Species-level niche-based studies have assumed that individual variation is either small or negligible compared to between-species variation (21). However, due to the strong spatial, temporal, and ontogenetic heterogeneity within populations, considerable individual niche variation is the norm rather than the exception (24, 26–28). Recent empirical evidence has repeatedly demonstrated the importance of individual variation in driving species’ coexistence (29, 30), ecosystem functioning (31, 32) and adaptive capacity (33).

Ignoring individual variation also reduces our capacity to make accurate predictions about the impacts of climate change on ecosystems (34–36). For example, if individuals and the populations they form are highly specialized to the local environments and differentiated across a species’ geographic range (37–39), a small change in mean environmental conditions can negatively impact all populations but may be judged unharmful by a species-level vulnerability analysis (e.g. 40). If individual niches are highly nested within one another (28), small environmental changes might have little impact on a species’ geographic range size but could extirpate specialized individuals that provide vital ecosystem functions (31, 41). Moreover, individual variation provides the fuel for evolutionary adaptation (42). Neglecting individual variation inevitably leads to biased estimation of niche dynamics and species distributions under climate change (43).

There are two main reasons for historically ignoring individual variation in niche-based studies. The first is operational: due to the scarcity of individual data, niche estimation has only been feasible at the species level for many studies that use opportunistic observational data (20). However, the tide has turned in recent years with the rapid accumulation of individual data such as those provided by animal GPS tracking (44, 45) camera trapping (46, 47). By far, the amount of individual animal movement data had exceeded the amount of all other biodiversity data combined (48–50). The rapid growth of individual data now provides a valuable opportunity to examine niche variation below the species level (26, 28) and better integrate biogeography with behavioral ecology (51). Broadly, it offers the potential for a more mechanism-driven prediction of large-scale biodiversity changes using a bottom-up approach (34, 52).

The second reason is conceptual. Ecologists have struggled to mathematically relate species niches to individual niches (22, 53, 54). The problem largely stems from the failure to reconcile different niche concepts over the years (17). The widely accepted Hutchinsonian niche concept, defined as the environmental conditions where a species has positive population growth rate, is usually infeasible to quantify empirically (2)(e.g., being too costly for large-scale analysis and until recently undefined at the individual level; (54)). As a result, ecologists are forced to use surrogate niche measures that bring inconsistency and confusion to the niche concept, such as habitat suitability (55) estimated from occurrence data (most commonly performed using species distribution models), putative functional differentiation calculated from species-level trait data (56), and thermal performance in physiological studies (57, 58).

Characterizing ecological niches from individuals requires a niche measure that is scalable for empirical studies across organismal levels and mathematically tractable for the development of theory. Despite the lack of consensus on what exactly is the meaning of the niche (16, 59, 60), an operational niche measure has emerged independently from several empirical applications and become commonly used and well-accepted - the niche as measured by the probabilistic distribution of a taxonomic unit in the environmental or trait space (61–63): in macroecology, the niche of a species is quantified by its relative occurrence rate in climate space (55); in community ecology, a species’ niche is often quantified by the distribution of individuals or biomass in the functional space (56, 64); in nutritional ecology, an animal’s dietary preference is measured by the relative frequency of food consumption in isotopic space (65, 66); in movement ecology, the niche of an individual is quantified by the utilization distribution or resource selection function in environmental space (67). This probabilistic niche measure not only benefits from a suite of increasingly mature multidimensional computational tools to describe their geometric features such as the volume and overlap (68, 69), but has also been used to develop ecological theories such as the theory of limiting similarity (70) and the trait-driver theory (64).

Previous use of the probabilistic niche measure to investigate individual niche specialization have been predominantly focused on the niche breadth (i.e. as measured by the variance of resource use) (21, 26). However, a more general approach to study individual variation of other important niche properties is strongly desired to better appreciate how species respond to climate change. One prominent example is the skewness of the thermal niche, as exemplified by the left-skewed thermal performance curve (71). The rapid decrease of thermal performance once past the optimal condition implies that individuals and species might respond strongly to warming once a certain threshold is crossed and result in sudden population declines (40, 72). To date, we have little knowledge about how individual variation drives the asymmetry of the species-level thermal niche.

In this paper, we propose a framework to relate estimated niches across organismal scales. Our primary goals are: 1) To show that the population- and species-level niches can be analytically derived from individual niches, which provides a convenient way to partition population-level niche properties (including but not limited to niche breadth and skewness) into individual contributions regardless of the underlying distributions. 2) To show that our framework is widely applicable to many ecological and evolutionary questions that require predicting species-level niche change from individuals, such as niche shifts under climate change. We demonstrate the utility of our framework using animal tracking data for two example species, the Gadwall (*Mareca strepera*) and the African bush elephant (*Loxodonta africana*), but our method is generalizable to any mobile organisms and other types of individual data.

## Methods

### Probabilistic niche measures

We briefly recap our definition of probabilistic measures of individual niches and population niches before elucidating their mathematical relationship.

The individual niche here is measured as the relative occurrence rate of an individual in the environmental space. In a one-dimensional example where the environmental variable is denoted by *x*, the niche of the *i*th individual is measured by the probabilistic density function, *f*(*x*)*_i_*, describing the individual’s relative occurrence rate in the environmental space. This measure follows the convention of how individual niche is quantified in movement ecology (26, 28).

The population niche is measured as the relative occurrence rate of individuals from the population in the environment. In the one-dimensional example, the population niche is measured by the probabilistic density function, *F*(*x*). This measure follows the logic of how population-level or species-level niches are quantified in species distribution modeling if individuals are randomly sampled from the landscape (55).

We have not explicitly distinguished from the fundamental niche and the realized niche in our treatment. Our framework is kept as general as possible to be applicable to both the realized niche and the fundamental niche as long as they are measured by probability density functions (73). Although in practice, we acknowledge that because the fundamental niche is difficult to estimate from observational data, the probability measure of the niche is most often applied to the realized niche.

The key to link population-level niches to individual niches is to realize that the probability of any individual from the population occurring in a certain environment can be calculated by a two-stage sampling process: first choosing an individual from the population, then choosing the environment for the individual to occur in. Let’s use a heuristic example to demonstrate this calculation.

### A heuristic example of the mixture distribution

For a heuristic example, let’s consider a population of a chameleon species consisting of three individuals that have distinctive temperature niches (Fig. 1A). The population niche can be derived using conditional probabilities: the relative occurrence rate of individuals from the population in environment *x* is calculated by the probability of choosing the *i*th individual in the population *w_i_*, multiplied by the relative occurrence rate of the *i*th individual in the environment *f*(*x*)*_i_*, then summed across all individuals in the population (Fig. 1B).

**Figure 1.**
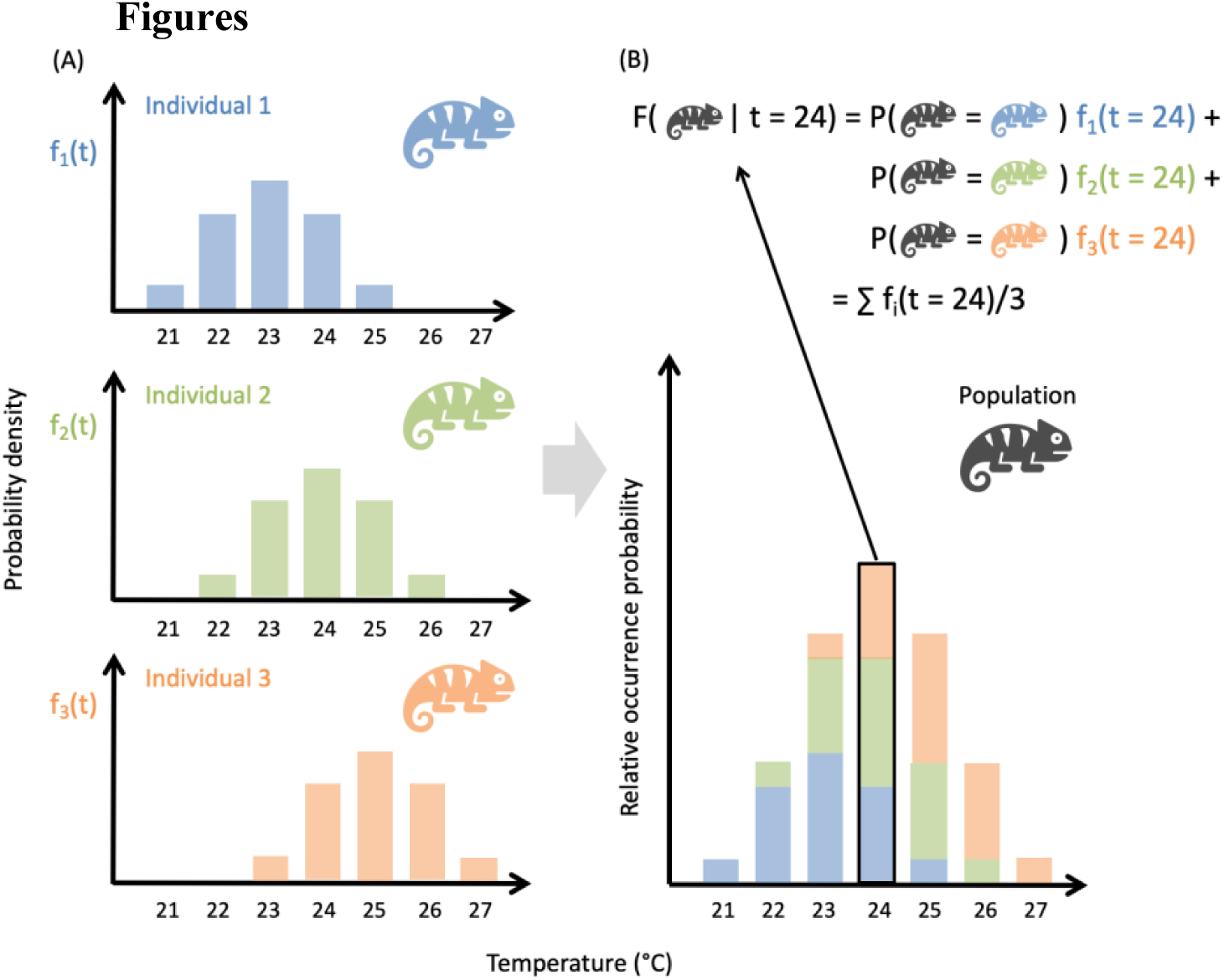
Conceptual figure of scaling from (A) individual niches to the (B) population niche. The population niche (B) is described by a mixture distribution of individual niches from three animals (A). f_1_(t), f_2_(t) and f_3_(t) represent the temperature niche of the three individuals.

Mathematically, the resulting population niche can be expressed as a mixture distribution of the individual niches, with *w_i_* representing individual weights (in this case, each individual has the same weight because they contribute equally to the population):

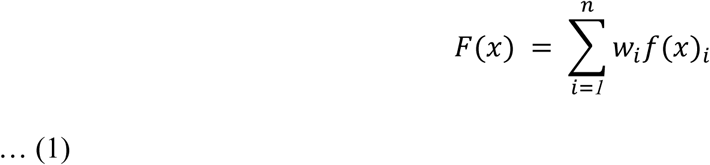

Alternatively, we can arrive at the same mixture distribution by converting an individual-level occurrence dataset to a species-level occurrence dataset by removing individual identity information (74, 75). For our heuristic chameleon example (Fig. 1), this procedure is equivalent to stacking the three individual niches together (Fig. 1A) and renormalizing it to get the probability density distribution that measures the population niche (Fig. 1B).

### Partitioning the population niche into its individual contributions

As well-established in the statistical literature, the moments of the mixture distribution can be derived from the moments of the component distributions (76). For a trivial example, the first moment of the population distribution (the population mean, μ) is just the weighted sum of the individual mean, μ_*i*_, where *i* denotes the *i* th individual and *w_i_* denotes the weight:

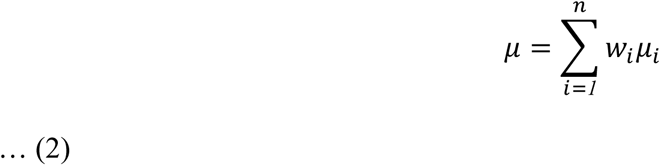

Nontrivial cases are shown by the individual partitioning of the higher moments:

Niche breadth is often represented by the second moment (the variance). In this case, the population niche breadth can be written as:

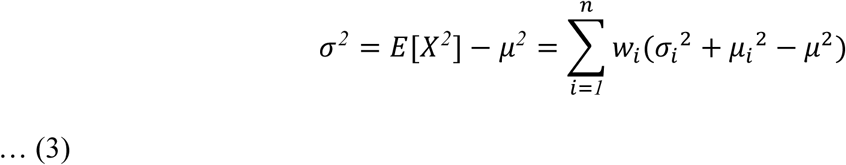

Where σ_*i*_*^2^* denotes the individual niche breadth. Equation 3 provides a natural partitioning of the population niche breadth into contribution through individual niche breadth, σ_*i*_*^2^*, and the contribution through niche position, μ_*i*_*^2^* − μ*^2^*. σ_*i*_*^2^* and μ_*i*_*^2^* − μ*^2^* can also be interpreted as the within-individual variation and between-individual variation in light of the analysis of variance. Note that to compare the relative contribution of the two components, μ_*i*_*^2^* − μ*^2^* must be non-negative to be interpretable. To achieve this, individual niche centers should be rescaled so that the population niche center μ = 0.

The third moment of the population niche (the skewness, γ) can be written as:

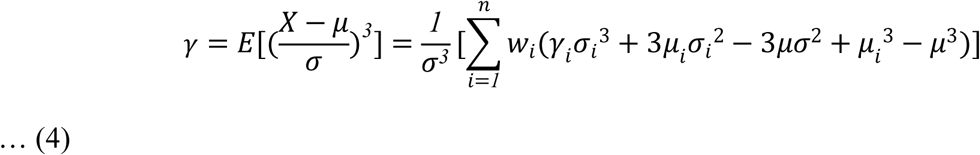

Where the population skewness is partitioned into a within-individual component γ_i_ σ_*i*_*^3^* and a between-individual component *3*μ_*i*_σ_*i*_*^2^* − *3*μσ*^2^*+ μ_*i*_*^3^* − μ*^3^*. The between-individual component is determined by the means and variances of individual niches, while the within-individual component is determined by the skewness and variances of individual niches. Alternatively, γ σ_*i*_*^3^* + *3*μ_*i*_σ_*i*_*^2^* − *3*μσ*^2^* roughly correspond to the contribution through niche breadth, and μ_*i*_*^3^* − μ*^3^* represent the contribution through niche position alone as in the case of variance partitioning.

### Multivariate niches

Equation 1 is equally applicable to the multidimensional niche space, although in this case an additive partitioning of all the moments is not applicable. Instead, we show that the ratio between the multivariate population niche breadth (as defined by the generalized variance) and the individual niche breadth can be partitioned into univariate components and a dimensionality component (Supplementary information, Appendix 1), and this ratio is closely related to Wilk’s Lambda in the multivariate analysis of variance (77).

### Beyond population niches

The implication of our framework goes beyond population niche estimates. In many ecological and evolutionary studies, the weights in equation 4 can be chosen to reflect the relative abundance in each group. Such a procedure can be used iteratively to build species-level niches from sub-populations (20), and to build clade-level niches such as genus-level niche and family-level niche (78) using species-level niche as the units with weights reflecting the number of species in each clade.

### Predicting future population niches

Our framework can also be used to make predictions about the change in population-level niches. For example, predation may disproportionately affect individuals with particular niche preferences. Individuals can also potentially reduce predation risks by having similar niches with each other (79, 80); another example is that climate change will change the fitness of individuals with different niches (81), and cause a population-level niche shift. In such cases we can modify the values of *w*_*i*_ such that they reflect the relative contributions of different individuals to the population in the future.

### Empirical example: niche shifts under climate change

For a simple empirical demonstration of our framework, we predicted the niche shifts of a Gadwall (*Mareca strepera*) population (82) (15 individuals, 2130 observations) and an African bush elephant (*Loxodonta africana*) population (83) (15 individuals, 99988 observations; Fig. S1) under an *ad hoc* climate change scenario and a naive dispersal assumption (that individuals are only allowed to move within the modeling domain). Detailed methodology of the analysis can be found in the supplementary information (S1).

Although more realistic assumptions can be made, we do not attempt to provide a rigorous climate change projection but to simply illustrate the predictive capacity of the presented framework.

Our analysis involves 4 steps:

1. Estimate individual niches (i.e. temperature selection functions) using the observed occurrence using GPS data and remotely sensed temperature data, with an inhomogeneous Poisson point process model (84).
2. For each individual, calculate the current and future climatic suitability for each 1km pixel over the landscape. The pixel level suitability is estimated from individual temperature selection functions.
3. For each individual, calculate the average suitability across the landscape (all 1km pixels) respectively for current and future climatic conditions. Then calculate the weights in equation 1 as the ratio between the average future suitability and the average current suitability. The higher an individual’s weight is, the more likely it will contribute to the future population niche.
4. Estimate the future population niche using equation 1 with the calculated weights, and compare it with the current population niche.

## Results

### Partitioning the population-niche: hypothetical scenarios

To conceptually demonstrate the utility of the partitioning framework, we consider three hypothetical scenarios made up of six individuals along a one-dimensional temperature niche space (Fig. 2). Each individual is assumed to have a normally distributed temperature niche. The ‘evenly distributed’ scenario assumes that individuals have the same niche breadth, and their niche position is uniformly distributed in the environmental space (Fig. 2A). The ‘clustered’ scenario assumes that individuals centered at the same niche positions have different niche breadth (Fig. 2B). The ‘nested’ scenario assumes that the individual niche position is correlated with the individual niche breadth (Fig. 2C). This requires the niches of the specialized individuals to be a subset of the niches of the generalized individuals.

**Figure 2.**
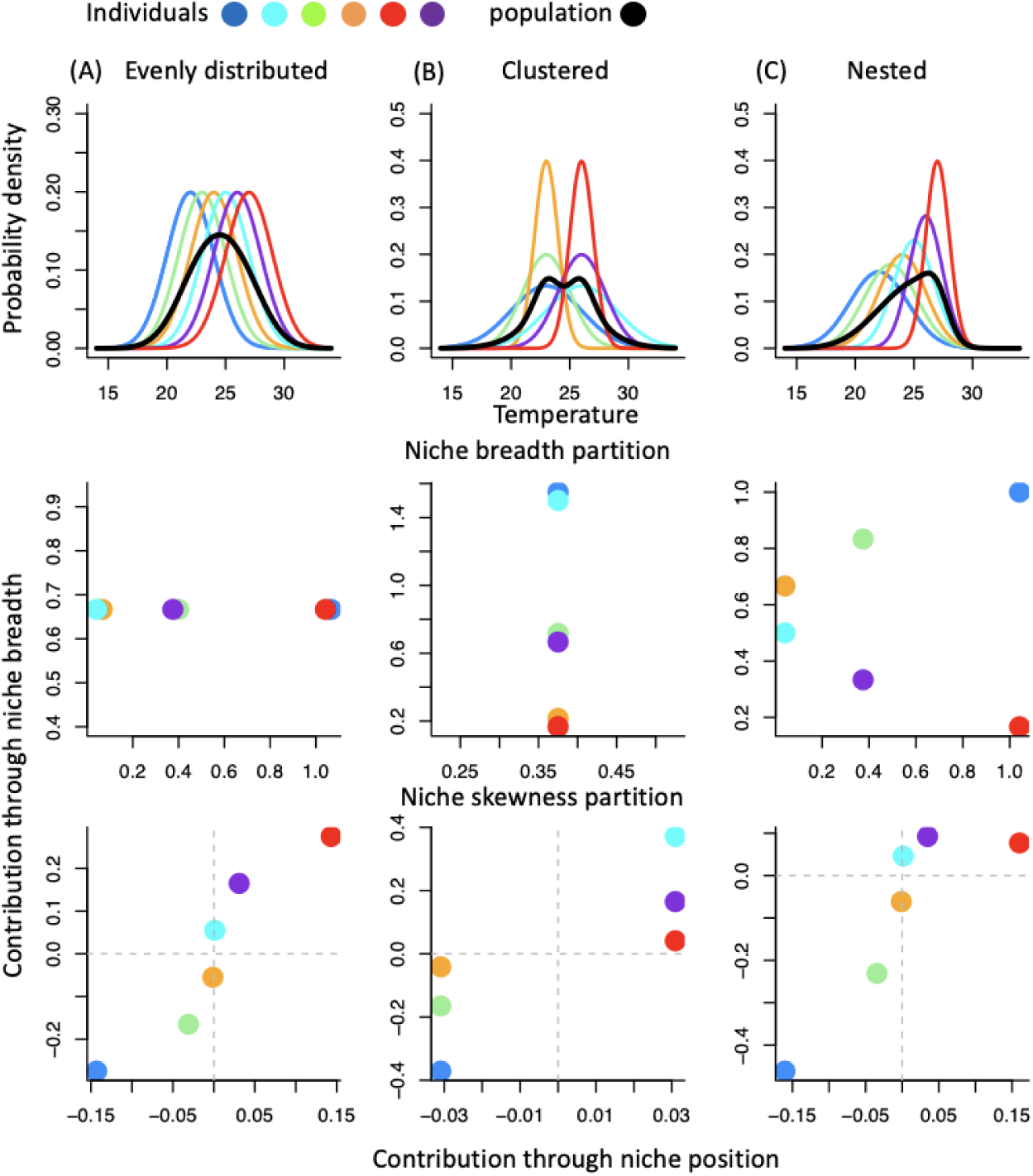
The conceptual figure for different scenarios of individual niche variations. Individuals are assumed to have a normally distributed environmental niche. Black lines represent the population niche, colored lines represent individual niches. (A) The ‘evenly distributed’ scenario assumes that individuals have the same niche breadth and evenly spaced niche center along a one-dimensional environmental axis. (B) The ‘clustered’ scenario assumes that individuals have different niche breadths and highly clustered niche positions. (C) The ‘nested’ scenario assumes that the niches of the more specialized individuals are a subset of the niches of the more generalized individuals. The second and third rows show the partitioning of the population level variance and skewness into individual contributions.

Under an even distribution of individual niches along an environmental gradient (Fig. 2A), the major variation in the individual contribution to the population niche breadth is through variation in individual niche positions, wherein individuals adapted to more extreme environments have a larger contribution to the population niche breadth than those adapted to intermediate environments. In contrast, if individual niches are highly clustered (Fig. 2B), then the major variation in the individual contribution to the population niche breadth is through variation in individual niche breadths, to which generalized individuals have a larger contribution to the population niche breadth. When the individual niches are nested, the bivariate plot of individual contribution through niche breadth and niche position shows a clear hump-shaped pattern (Fig. 2C). Furthermore, the ‘nested’ scenario also shows that a skewed population-level niche can arise from the clustering of specialized individuals in extreme environments even when individual niches are symmetric and have no skews.

### Empirical examples: niche shifts under climate change

The Gadwall population (Fig. 3, top row) is characterized by substantial individual niche variations (Fig. 3A). Overall, 66.9% of the population niche breadth (3.37) is contributed by individual niche breadth rather than variation in individual niche position. The interaction between individual niche breadth, position, and skewness (eqn. 4; corresponding to the contribution through niche breadth in skewness partitioning) contributes strongly (65.0%) to a positive-skewed population niche (0.80), while the effect of individual skewness alone is negligible (−0.0075).

**Figure 3.**
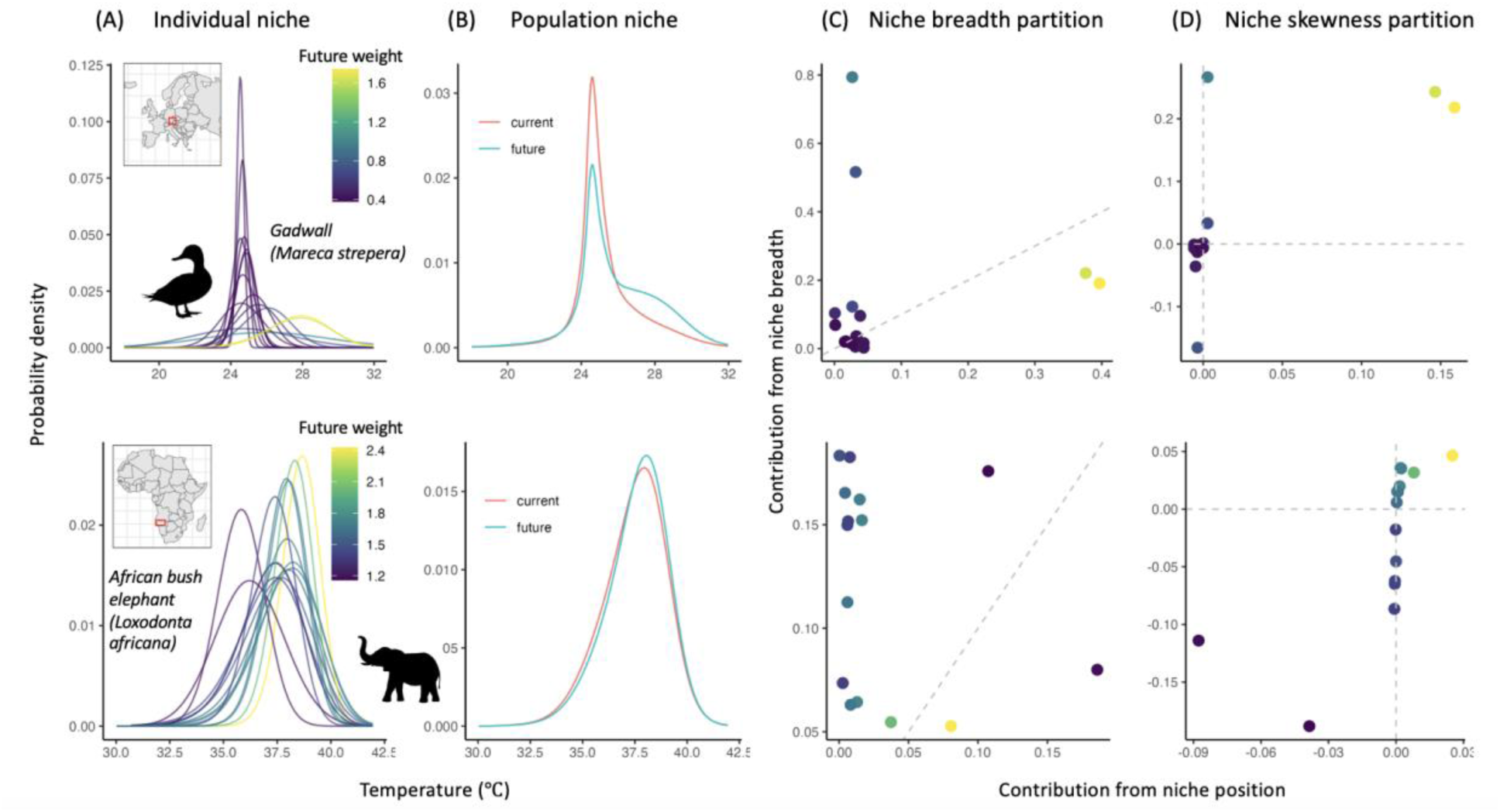
Individual niches (A) and population niches (B) under future warming of 2.4 °C in 15 Gadwall from southern Germany (top panels) and 15 African bush elephants from the Etosha National Park (bottom panels). (C) (D) respectively show the partitioning of population niche variance and skewness into individual contributions. Colors in (A) (C) (D) refer to future weights of individual niches in the warming scenario. The weight of each individual is calculated by the ratio between future habitat suitability and current habitat suitability. The grey dashed lines in (C) show the 1:1 lines. The horizontal and vertical grey dashed lines in (D) mark the 0 lines.

Under a hypothetical scenario of 2.4 degrees of warming (∼ RCP 3.4 climate scenario), the future population niche (Fig. 3B) is estimated to shift towards a higher mean (from 25.37 to 25.98 °C), a wider niche breadth (from 3.37 to 4.97), and to be less skewed (from 0.80 to 0.28). A closer examination of the individual partitioning of the population niche breadth and skewness shows that the shift towards wider niche breadth is predominantly driven by a strong selection for warm-adapted or generalist individuals (Fig. 3C), while the change of skewness is driven by the loss of thermal specialists that clustered at the relatively cold end of the population niche (Fig. 3D).

The case of the African bush elephants provides an interesting contrast to the Gadwall example (Fig. 3, bottom row). The individual niche breadth of the African bush elephants contributes even more to the population niche breadth (79%). In terms of the skewness, the individual niche skewness contributes to 51% of the strongly negative skewed population niche (−0.46).

As a consequence of the predominant role of the within-individual niche contribution in driving the population niche (Fig. 3A). The 2.4 degrees of warming causes a much lower magnitude of niche shift at the population level (Fig. 3B), with only a 0.1 °C shift of the niche center (37.5 to 37.6 °C), a slight decrease of niche breadth (2.33 to 2.21), and increase of the magnitude of skewness (−0.46 to −0.53). The individual partitioning of the population niche shows that although the selection for generalists (Fig. 3C) and for highly negatively skewed individuals (Fig. 3D) exists, it is unlikely to cause a drastic change in population niche due to the lack of between-individual variation.

In terms of spatial prediction, the Gadwalls are estimated to shift towards higher elevations in face of warming (Fig. S1A). Our individual-based model predicts a less concentrated preference in the geographic space (Fig. 4A) compared to the conventional species-level model (Fig. 4B), and higher suitability at low elevation areas (Fig. 4C) largely due to an increase in future niche breadth (Fig. 3B). The two models provide discrepant spatial predictions (*R*^2^ = 0.85), especially towards the more suitable pixels (Fig. 4D). The African bush elephants are predicted to shift towards inland areas under warming (Fig. S1B). Because little niche shift is incurred by warming (Fig. 3B), the individual-based model and the conventional species-level model yield almost identical spatial predictions of habitat suitability (*R*^2^ = 0.98; Fig. 4D).

**Figure 4.**
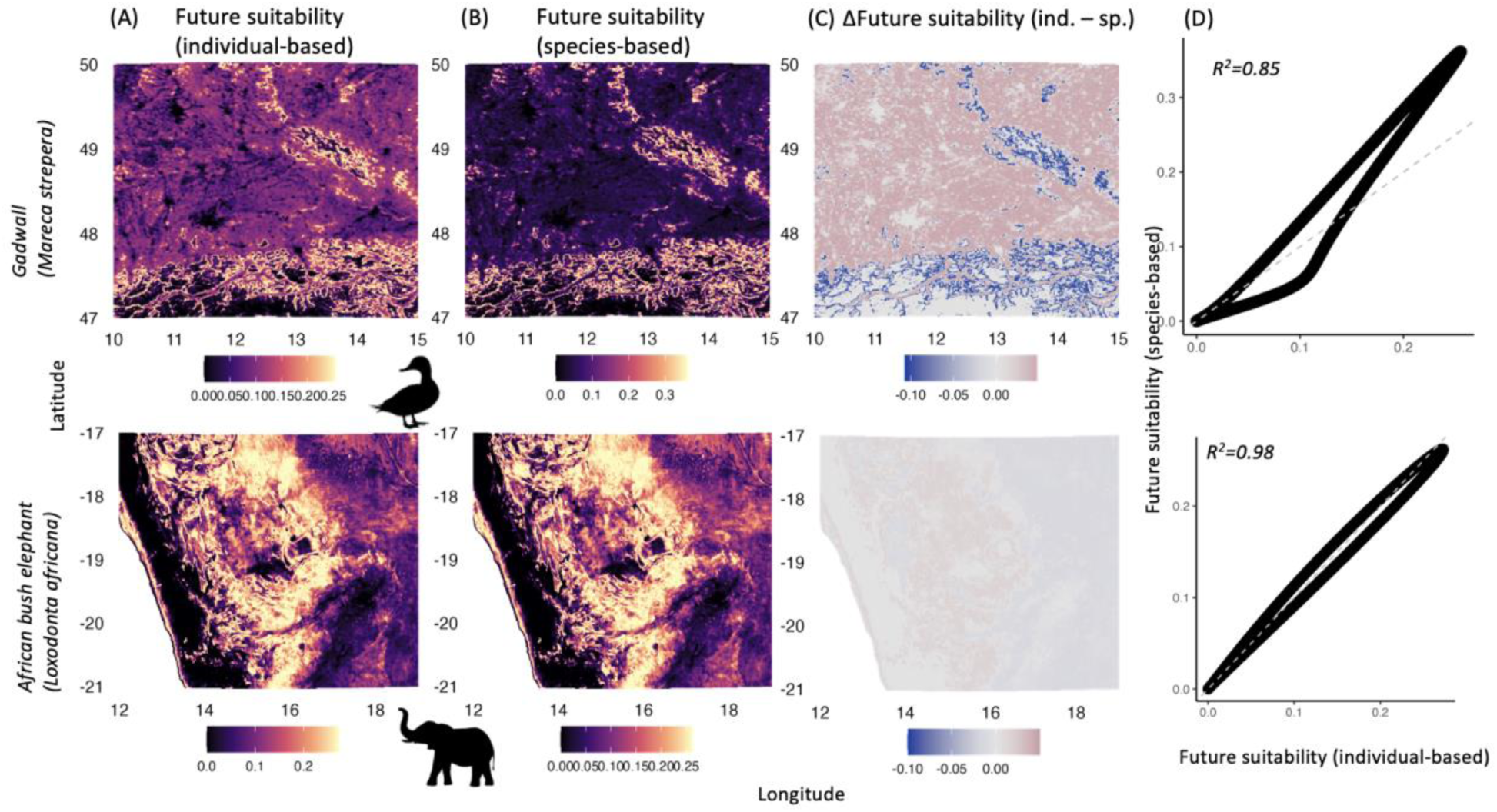
(A) The future suitability estimated from individual-based niche models (accounting for future niche shift). (B) The future suitability estimated from species-based niche models (not accounting for future niche shift). (C) Difference between future suitability estimated from individual-based model and species-based model. (D) Scatter plot of future suitability estimated from species-based model against future suitability estimated from individual-based model. The grey dashed lines show the 1:1 line.

## Discussion

In this study, we introduce a bottom-up approach to characterizing population-level and species-level niches from individual niches. We demonstrate the utility of this framework for partitioning individual niche contributions and predicting species-level niche shifts. We do not characterize the species-level niche from individual-level process in the *sensu stricto* Hutchinsonian definition, as attempted by recent studies (53, 54). Rather, we adopt a more operational measure of niche, the probabilistic occurrence niche, that stems from the use of species distribution modeling in macroecology (63) and resource utilization function (67) in movement ecology. By following the logic of how niches are characterized in these disciplines, we show that there is an elegant mathematical link between individual-level niche and higher organismal-level niches that opens many possibilities for future theoretical and empirical studies to investigate ecological and evolutionary processes across scales.

### Implications for intra-specific niche variation

The individual niche partitioning enabled by our framework recovers the traditional top-down variance partitioning approach to study within-individual and between-individual niche variation (21, 85), but further ties the contribution to species-level niche formation to specific individuals. Moreover, our method goes beyond investigating the niche breadth (as measured by the variance) and extends to all moments of the niche such as skewness, regardless of the underlying distributions.

One prominent example of the importance of considering skewness in species niche is the use of thermal performance curves (TPC) in assessing the risk of climate change (71). Thermal performance curves are usually derived from individual-based physiological experiments (86, 87). The generally highly left-skewed individual-level TPC predicts that organisms will respond to warming more abruptly than cooling (72). One key question of applying experimentally-derived thermal performance curves to predict species’ distribution at the macroecological scale is how well individual-level TPCs represent species-level thermal response (71). Our study shows that the species-level skewness of a climatic niche is not only determined by individual-level skewness but also determined by the interaction between individual-level niche position and niche breadth. As clearly demonstrated in our empirical examples, while the population skewness of the African bush elephants is largely determined by individual-level skewness, the Gadwall population shows the opposite pattern. To what extent does individual-level niche skewness dictate species-level skewness in general presents an interesting arena for future empirical research.

The ability to disentangle and characterize higher-level niches at the individual level enables us to account for life-history specific, spatial and temporal variation of species. For example, some bat species have highly segregated sex-specific habitat requirements (88), which can be incorporated under our framework to improve species-level niche estimates. In addition, accounting for population demographic structure will allow a better understanding about how species-level niche changes over time. Larval frogs (tadpoles) typically have distinct habitat, dietary and thermal requirements than adults (89). During certain parts of the year, tadpoles can greatly outnumber adults, and in other parts of the year they are completely absent, which can heavily sway population-level niche estimates if such temporal variation of demographic structure is not accounted for. Our framework can also be used to combine different temporal niches to obtain a species’ annual niche response. For example, our findings suggest that migrating bird species that have different niche tracking strategies over seasons (90) could have drastically different mean annual niche responses: species that have more variable seasonal niche breadths but fixed niche centers are more likely to have a highly skewed mean annual niche. In summary, our framework helps better understand the source of intra-specific niche variation and allows us to make more effective conservation decisions for different species depending on their biological characteristics.

### Implication for species distribution models

Our method also provides a natural way to build species distribution models (SDM) from individual resource selection functions (52, 74). The conventional way of accounting for individual-level variation in SDMs typically treats individual variation as a random effect in a generalized mixed effect model (52), and uses the mean regression coefficients to represent the population-level response. In our empirical examples, we show that to characterize species-level SDM from individual data that is comparable to the occurrence-derived SDMs (74), the population response should be calculated as a mixture distribution of individual resource selection functions rather than using their mean regression coefficients. The difference between the two approaches of modeling population-level or species-level response is due to Jensen’s inequality (the mean response is not equivalent to the response of the mean; (91)). An important limitation of our method is that it depends strongly on the random sampling of individuals across space and time. The standard practice of removing duplicate records in a grid cell (using grid cell rather than individual as the sampling unit) before fitting a SDM (55) might hinder the integration of our approach with traditional methods. However, we expect that the gap will be increasingly bridged by the rapid growth of individual data (45, 49). To fully tap the potential of our framework for SDM, further investigation on the optimal sampling of individuals that yield the highest gain of information regarding habitat requirement (92, 93) is also needed.

### Implication for climate change vulnerability assessment

A framework that can utilize individual-level information is especially important for climate change research. First, individual-based niche models using movement data may reshape estimates of animal redistributions and more accurately account for the effect of dispersal under climate change (94, 95). Second, animals may evolve their thermal preferences to adapt to local conditions (96), which may occur at different rates based on body size and other individual-level factors (97, 98). Third, given that extreme weather events are expected to increase in severity and frequency under climate change (99), incorporating shorter term climatic anomalies in niche models could determine whether the exposure of individuals to extreme climatic conditions goes beyond their niche limits. Importantly, the thermal limits of individuals are expectedly narrower than those of the species as a whole, suggesting that individual-level climate change vulnerability may be masked by species-level models. Moreover, our findings suggest that different individual-level niche configurations are likely to produce different species-level responses to climate change. For example, in the ‘clustered’ scenario (Fig. 2), populations are more likely to undergo an abrupt collapse of half of their individuals under climate change; in the ‘nested’ scenario, the niche structure might have the potential to alleviate the threat of climate change not only by favoring warm-adapted individuals but also generalists in the future (Fig. 3), as illustrated in the Gadwall example. Without a thorough understanding of individual niche variation within species, it is impossible to accurately forecast species’ extinction risk to climate change.

### Implications for ecological theory

The utility of the framework goes beyond empirically partitioning the niche or predicting niche shift and lies in its potential for developing individual-based ecological theories. Previous theoretical studies have examined how the evolution of niche breadth is influenced by both intra- and inter-specific competition, abundance of resources and the strength of natural selection (100, 101). Our framework enables the investigation of the general shape of the ecological niche (102) which includes important attributes such as skewness.

Another important field that can clearly benefit from our findings is the investigation of functional traits. Traits are often used as proxies for niches, in which cases, the *f*(x)*_i_* in equation 2 can be used to describe the distribution of biomass or abundance of species in the functional trait space. The partitioning of trait attributes such as skewness and kurtosis into within-species and between-species components fits nicely into the regime of the trait driver theory, which links the distribution of traits to ecosystem functioning, productivity, stability and climate change (64, 103).

### Connection to the Hutchinsonian niche

The link of our approach to the *sensu stricto* Hutchinsonian niche is not through the quantification of environment use *per se*, but through the weight function in equation 1. In our demonstration of the niche shifts examples (Fig. 3), we have implicitly assumed that the future environmental suitability of an individual represents its relative fitness under climate change. To further accommodate the Hutchinsonian niche concept, we can use absolute fitness rather than relative fitness to calculate weight in equation 1. If fitness information on individuals can be obtained independently from occurrence data, then we can select or weight the contributions of individuals to population-level Hutchinsionian niche based on their contributions to the long-term persistence of the species (54). For example, consider the case that inferior competitors of an animal population leaving high-quality natal habitats for inferior habitats, and many of these animals will likely die or may not reproduce given poor habitat quality. If we were to estimate the species niche for all individuals, we may estimate a wider species niche than the actual Hutchinsonian niche, as the peripheries of this estimated distribution do not contribute to species’ persistence. This problem of overestimating the Hutchinsonian niche with observed occurrence is particularly insidious in the case of ecological traps: individuals may actually be attracted to habitats that actually harm individual fitness (104), leading to over-estimation of the species-level reliance on poor habitats.

## Conclusions

The unification of niche estimates across organismal levels provides a valuable opportunity to further advance ecological theories, refine climate change predictions, and inform biodiversity conservation efforts. The increasing integration of individual-level studies into global biodiversity research will greatly enrich our understanding of biodiversity changes across spatial, temporal and organismal scales.

## Data availability

The data used in this publication were accessed from publicly available sources. The gadwall data set can be found at: https://doi.org/10.5441/001/1.26dg08hv and the elephant dataset can be found at: https://doi.org/10.5441/001/1.3nj3qj45. The datasets to reproduce the study will be stored in Dryad.

## Code availability

Computational scripts to replicate the study are available on Github (https://github.com/lvmuyang/ind2pop-niche) and will be stored in Zenodo upon acceptance of the manuscript.

## Acknowledgements

We thank Jetz lab members for discussion and comments on the manuscript.

## Author contributions

M.L. conceived the study, derived the theoretical results and led the writing. S.Y. and M.L. performed the empirical analysis. All authors contributed to writing the manuscript.

## Supplementary information. S1

### Partitioning the moments of the mixture distribution

For a mixture distribution (with the random variable denoted by X) with probability density function defined by:

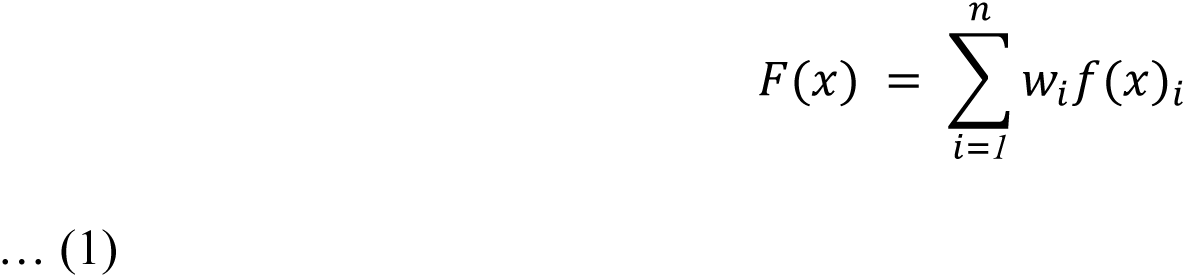

Its moments can be derived as follows:

Firstly, for any function of *x*, *q*(*x*), the expectation of *q*(*x*) is the weighted sum of the expectation of *q*(*x*)*_i_*:

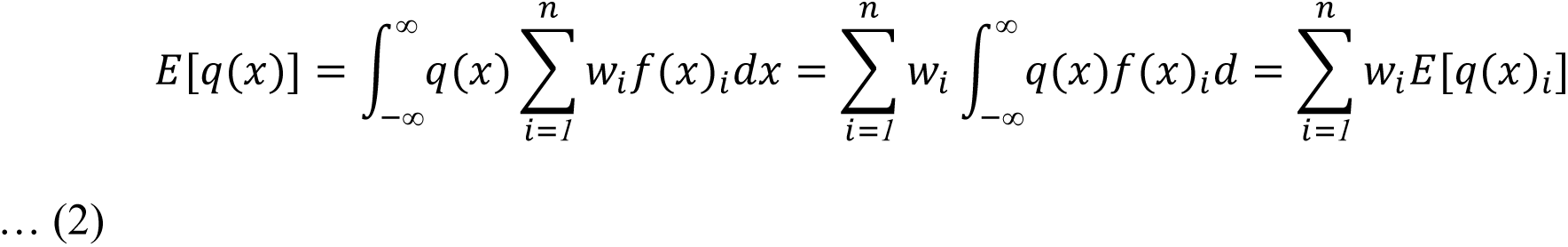

Then it is well known in the statistical literature that the moments of the mixture distribution can be deduced from the moments of the component distributions:

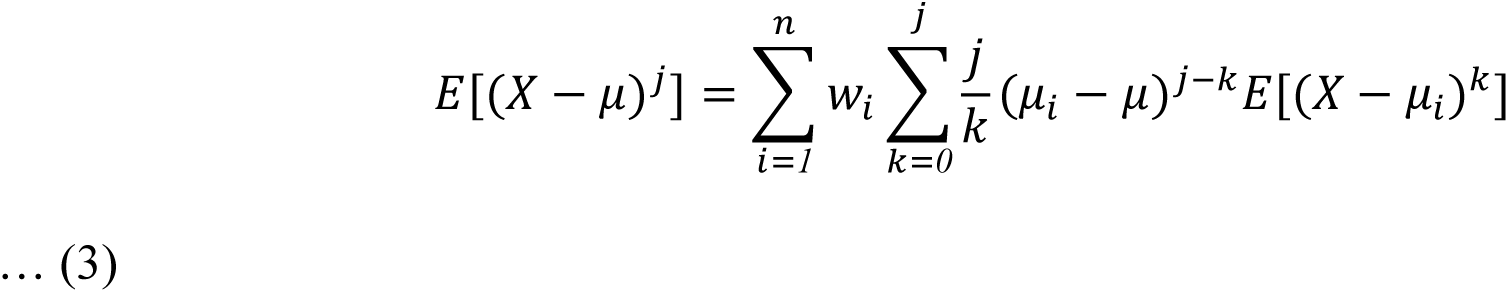

Where μ is the mean of population niche, μ_*i*_ is the mean of the individual niche *i*, and *j* denotes the *j*th moment.

### The partitioning of the multivariate niche breadth

For the multivariate environmental niche, equation 1 can be generalized as:

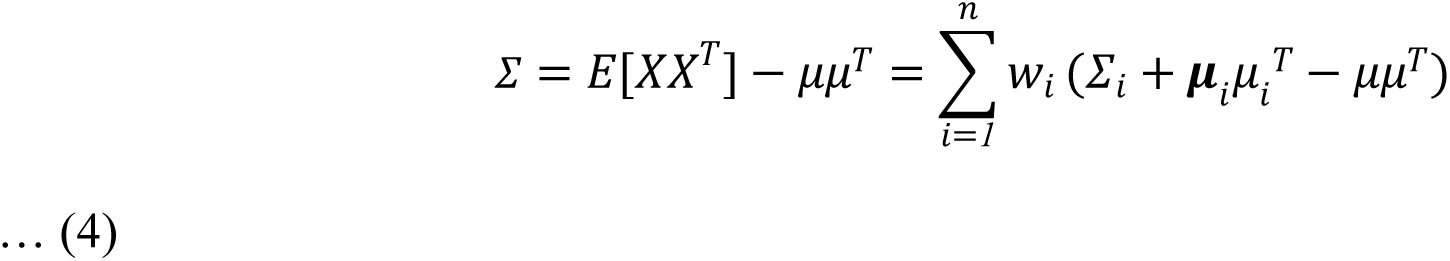

Where *X* = [*X_1_*, *X_2_*…*X*_*p*_] and μ = [μ*_1_*, μ*_2_*…μ_*p*_] are the vectors of the *p* environmental variables and the mean. Σ is the covariance matrix, and the multivariate niche breadth can be measured by the generalized variance |Σ| (the determinant of the covariance matrix).

The ratio between mean individual niche breadth and population niche breadth is equivalent to the Wilk’s lambda statistic in the multivariate analysis of variance, which can be further partitioned into univariate niche breadth ratios and the ratio of niche dimensionality:

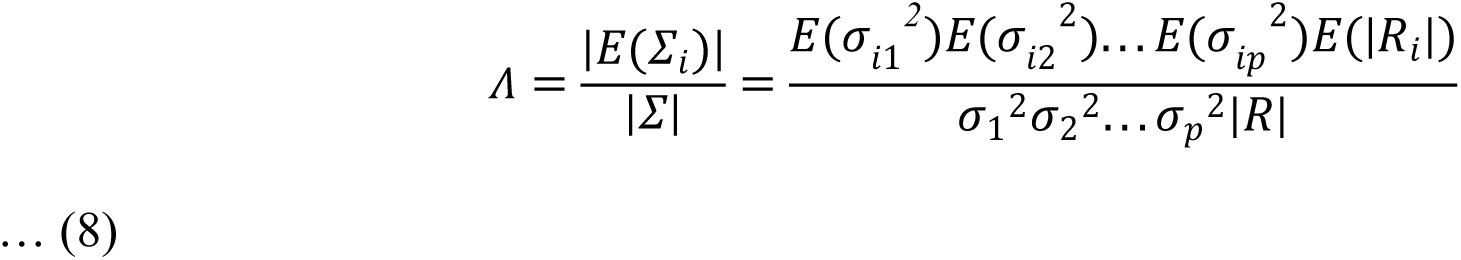

Where *R*_i_ and *R* are the correlation matrix of individual niche i and the correlation matrix of the population niche.

### Detailed methodology for the empirical individual niche analysis example

We gathered existing tracking data for 15 Gadwalls tracked in southern Germany in 2009 and 2010 and 15 African bush elephants tracked in Etosha National Park between 2008 and 2014. We used the MODIS Land Surface Temperature (LST) at 1km spatial resolution (MOD11A1) (https://developers.google.com/earth-engine/datasets/catalog/MODIS_061_MOD11A1) for the respective sampling period for the two species to perform the niche analysis.

We restricted Gadwall observations to summer months only (Julian days 121 to 243) and only retained individuals with a tracking duration of at least 30 days. We restrict our analysis to the modeling extent containing the minimum convex polygon surrounding all observations of Gadwall individuals plus an *ad hoc* 5 km buffer to accommodate a moderate amount of future dispersal from current locations. There are in total 2130 observations for the 15 Gadwall individuals and 99988 observations for the 15 African bush elephants individuals used for subsequent analysis.

**Figure S1.**
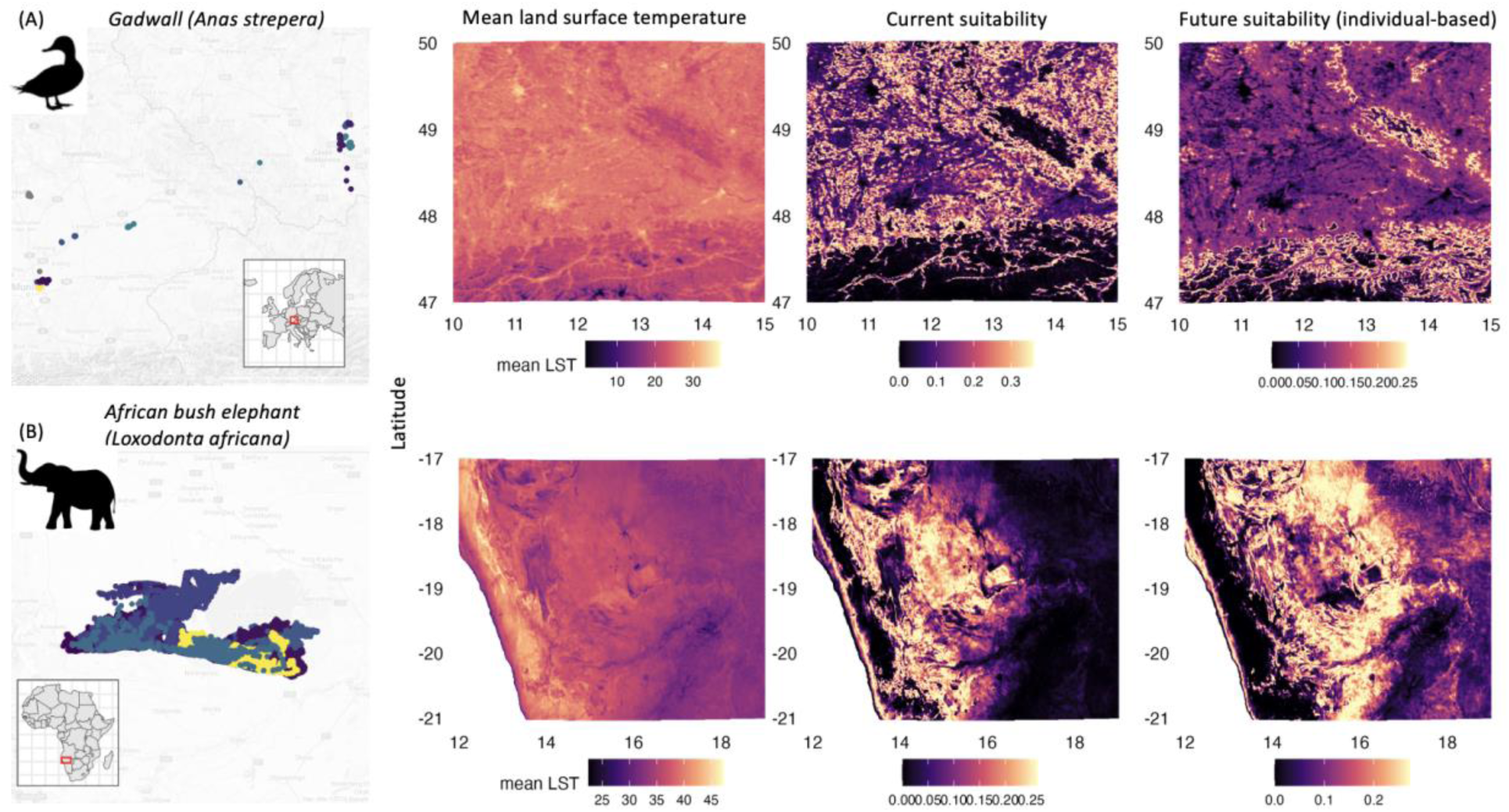
(A) GPS-tracked observations for the Gadwall and African bush elephant populations; (B) Mean land surface temperature during the tracking period. (C) Current population-level suitability. (D)Future population-level suitability estimated from our individual-based model.

Because of the temporal auto-correlation of the tracking data, estimation of the temperature selection functions directly through annotated data will be biased. To overcome this issue, we first estimate the home ranges of individuals using the autocorrelated kernel density estimator (akde) with the ‘ctmm’ package (Fleming et al. 2015). We then extracted the mean LST values within the 95% quantile of the ‘akde’ home range size for estimating the temperature selection functions.

We first resampled 10000 points within the home range using the probability density estimated from ‘akde’ as weight to generate presence-only points. We then sampled 100000 points from the whole modeling domain to obtain the background distribution. The temperature selection function was estimated through an inhomogeneous Poisson process model (Renner et al. 2015). Polynomial terms of the temperature up to the third order are included as predicting variables to account for skewness of the distribution.

Note that an alternative approach is to estimate temperature selection with the step selection function (Michelot et al. 2024), which is not attempted here for the simplicity of the demonstration.

We calculated the current suitability using the mean LST layer (restricted to the same period of interest as the individual data) and calculated the future suitability by adding the 2.4 degrees of warming (which approximates an RCP 3.4 climate scenario) to the current temperature. Future population suitability is calculated by first estimating future population niche using equation 1 with weights reflecting the relative fitness of individuals under climate change, then applying the future niche model to the warmed landscape. To compare it with conventional species-level niche models without niche dynamics, we also calculated the future population suitability with the current population niche using equation 1 with equal weights. The predictions of the two different niche models are plotted against each other for visualizing their discrepancy.

